# The over-activated peritoneal immune environment in endometriosis is characterised by a lack of PD-1 inhibition

**DOI:** 10.1101/2024.08.30.610573

**Authors:** Anna Tresso, Niharika Thota, Chloe James, Natasha Borash, Emily Brennan, Shima Bayat, Beverley Vollenhoven, Thomas Tapmeier

**Affiliations:** Department of Obstetrics and Gynaecology, Monash University, Clayton, Australia; The Ritchie Centre, Hudson Institute of Medical Research, Clayton, Australia; Department of Cellular, Computational and Integrative Biology, University of Trento, Trento, Italy; Women and Newborn’s Program, Monash Health, Clayton, Australia

**Keywords:** Endometriosis, full-spectrum flow cytometry, inflammation, peritoneal fluid, innate immunity, adaptive immunity, CD69^+^ T cells, PD-1, immune checkpoints

## Abstract

**Background:** Endometriosis is characterised by chronic inflammation in the peritoneal cavity causing acute and chronic pelvic pain, largely explained by dysregulation in the immune environment within peritoneal fluid. The activation status of the peritoneal immune cells is unclear. In addition, a comparison with the status of the systemic immune system is desirable to explore avenues of diagnosis and treatment of endometriosis-related inflammation and pain.

**Objective(s):** To investigate the immune environment in endometriosis in peritoneal fluid and blood by full-spectrum flow cytometry with a focus on activation and inhibition of immune cells.

**Study Design:** This was an observational study in patients undergoing laparoscopy for diagnosis or treatment of peritoneal endometriosis or for unrelated conditions; PF was collected from n=5 endometriosis patients and n=4 controls, blood from n=4 endometriosis patients and n=3 controls. Data were analysed for statistical significance using ANOVA, the Kruskal-Wallis or Mann-Whitney U test, with a p-value below 0.05 considered significant.

**Results:** We observed a prevailing of myeloid immune cells in the peritoneal fluid as opposed to lymphoid cells in the blood. The main differences between endometriosis and control samples, however, were found in the smaller compartments, i.e., in lymphoid populations in peritoneal fluid and myeloid populations in blood. PD-1 levels in peritoneal fluid endometriosis samples were significantly lower than in controls (p<0.05).

**Conclusion(s):** The immune checkpoint PD-1 could be a new angle of treating endometriosis-related inflammation and pain in women suffering from this chronic and intractable condition.

**Tweetable:** Lack of PD-1 plays a role in endometriosis-related inflammation.

**AJOG at a glance:**

- The inflammatoru immune environment in endometriosis needs investigating as it iis causative of the pain, i.e., the predominant symptom.
- We found a lack of PD-1 expression on peritoneal fluid cells in endometriosis compared to controls.
- This could explain the persistent inflammation and open avenues of treatment.

---

Endometriosis is an oestrogen-dependent chronic inflammatory disorder characterised by chronic and acute pelvic pain, dysmenorrhea, pain at ovulation, dyspareunia, abnormal bleeding, pain with bowel movements, fatigue, and infertility^1,2^. Treatment options are limited, with non-steroidal anti-inflammatory drugs (NSAIDs), acetaminophen, or opioids used to manage endometriosis-associated pain; hormonal treatments like combined hormonal contraceptives, progestogens, gonadotropin releasing hormone (GnRH) agonists or antagonists used to shut down the menstrual cycle and thus alleviate endometriosis-related symptoms^3^; and surgery to try and remove endometriosis lesions^4^. However, the latter option does not guarantee the prevention of disease recurrence and pain^5^.

Endometriosis-related pain is thought to be triggered by ‘extra-uterine menstruation’ from lesions^6^, and a deficiency in the immune response to the presence of disseminating endometrial debris the reason for lesions persisting in the first place^7^. A connection between endometriosis and autoimmunity has long been suspected^8^. However, the role of the innate and adaptive immune systems in endometriosis is unclear^9^. Decisive roles in endometriosis pathophysiology have been ascribed to the myeloid compartment, with macrophages responsible for promoting or preventing disease progression depending on their location or origin^10^. Overall, there is evidence of M1 macrophage predominance in the eutopic endometrium, while M2 macrophages (Mϕ) are more abundant PF of endometriosis patients^11,12^.

However, in the lymphoid compartment, T cells are thought to orchestrate a (deficient) immune response in peritoneal fluid^13^. Interleukin (IL-)8 induces cell adherence to extracellular proteins, while angiogenic factors like vascular endothelial growth factor (VEGF) support blood supply^14^. Chronic inflammation and cytokine production are further enhanced by overexpression of nuclear factor κB (NFκB), production of reactive oxygen species, expression of IL-17, and activation of mitogen-associated kinase (MAPK) signalling pathways^15,16^. These attract monocytes, eosinophils, and T lymphocytes, further contributing to the inflammatory response associated with endometriosis^17^. NK cell cytotoxicity has been found deficient in endometriosis patients^18–20^, while Dendritic Cells (DCs) may on the one hand attenuate lesion development through activation of T lymphocytes, on the other hand the presence of endogenous DCs is necessary for lesion formation^21,22^. Regulatory T cells (T_REG_) have a role in endometriosis pathogenesis by promoting immune suppression and limiting the immunological response to endometriotic lesions^23,24^. T helper 2 (T_H_2) and T helper 17 (T_H_17) cells are overrepresented in endometriosis^23^. A subpopulation of CD69^+^ T cells was found increased in the peritoneal fluid with reduced expression of markers associated with T cell function; this may be less functionally active than its counterpart in blood^25^.

To investigate the connection between all of these immune cell populations and the disease, high-dimensional flow cytometry data have become the tool of choice. Full-spectrum flow cytometry, FSFC enables the development of high multiparametric panels, which allow the identification of a larger number of markers per cell and can provide more comprehensive information about the immune system than conventional flow cytometry or even mass cytometry^26,27^.

Here, we hypothesize that aberrant activation of inhibitory T cells such as FOXP3+ T_REG_ or PD-1+ CD8+ T cells lie behind the persistence of endometriosis lesions. We therefore aim to characterise the cellular immune environment including differences in immune checkpoint activation and the overall activation status of T cell populations.

## Material and Methods

### Participants

All samples from consenting women were collected under Human research Ethics Committee approvals 01067B with 20-0000-882A or under 20-0000-159A. The participants (Supplementary Table 1) were women between 18- and 45 years old who were undergoing laparoscopy for suspected endometriosis or unrelated health conditions. After obtaining informed consent, peritoneal fluid (PF) and peripheral blood (PB) were collected at the time of surgery. Exclusion criteria were pregnancy, malignancy, and menopause. Hospital surgical records were used to confirm the diagnosis of endometriosis and the disease stage, hormone state, and menstrual cycle stage.

### Samples

PF samples were collected in 50 mL Falcon tubes and processed between 1-6 hours after collection. Briefly, PF was centrifuged at 3000 rpm for 5 minutes at RT for supernatant removal. If the PF showed RBC contamination, a RBC lysis step was performed, otherwise the cell pellet was washed in PBS and centrifuged at 500 g for 5 min at RT before being resuspended in PBS for cell counting. Blood samples were processed as described earlier^28^. Briefly, the cells were pelleted at 2000 g for 5 minutes at RT and the supernatant removed. The RBCs in the pellet were lysed using RBC lysis buffer (Cell Signalling, NEB) in three sequential lysis steps in separate falcon tubes, then reunited, washed in PBS and counted.

### Full Spectrum Flow Cytometry (FSFC)

Antibodies used are listed in Supplementary Table 2. All antibodies were titrated for optimal staining. For measurements, 2 x 10^5^ cells in stain buffer (2% FBS, 0.5% EDTA in 1X PBS) were labelled with single antibodies and 1 x 10^6^ cells with the full panel according to the manufacturer’s instructions. 2.5 μL of working solution ViaDye Red Fixable Viability Dye was added to the cell pellet. Tubes were centrifuged at 400 g for 5 minutes at RT, followed by supernatant decanting and blotting on a paper towel. After a 20-minute incubation step in the dark, samples were washed in 3 mL of Stain Buffer and centrifuged at 400 g for 5 min at RT followed by decanting and blotting on a paper towel. Each sample was resuspended in 150 μL of Stain Buffer. For intracellular staining, tubes were then fixed in 300 μL of 1% paraformaldehyde (PFA), permeabilized in 0.1 X Triton X-100 in deionized H_2_O and stained with anti-FOXP3. Cells were washed in stain buffer and resuspended in 300 μL of stain buffer for acquisition (methods adapted from DOC-00504 Rev. A, Cytek Biosciences, 2023^29^ and Kit DOC-00517 Rev. A, Cytek Biosciences, 2023 ^30^).

### Sample acquisition

All samples were measured in the Cytek Aurora (configuration 5L UV/V/B/YG/R) using SpectroFlo® software. Daily QC was performed and passed with SpectroFlo QC beads before sample acquisition. The same acquisition settings were used for each sample type. Gating scheme in Supplementary Figure 1. Autofluorescence was compensated for using unstained samples for each method of preparation. Single stain controls were established on PBCs, PFCs, and CompBeads for accurate unmixing. Cell populations were defined by markers as laid out in Supplementary Table 3.

## Statistical analysis

All data and plots were analysed with GraphPad Prism using two-way ANOVA, the non-parametric Kruskal-Wallis test or the Mann Whitney U-test. Asterisks indicate the statistical significance. *p < 0.05.

## Results

Our data showed the peritoneal fluid cells to be largely of the myeloid compartment (Fig. 1, A) and the blood cells dominated by the lymphoid compartment (Fig. 1, B). Interestingly, some population seemed exclusively present in one or the other sample species, while some overlap was also observed. This indicates tight regulation of the transition of cells from the circulation into the peritoneal cavity, and vice versa. The visualisation approach effectively pinpointed differences between cell populations in patients and controls.

**Figure 1:**
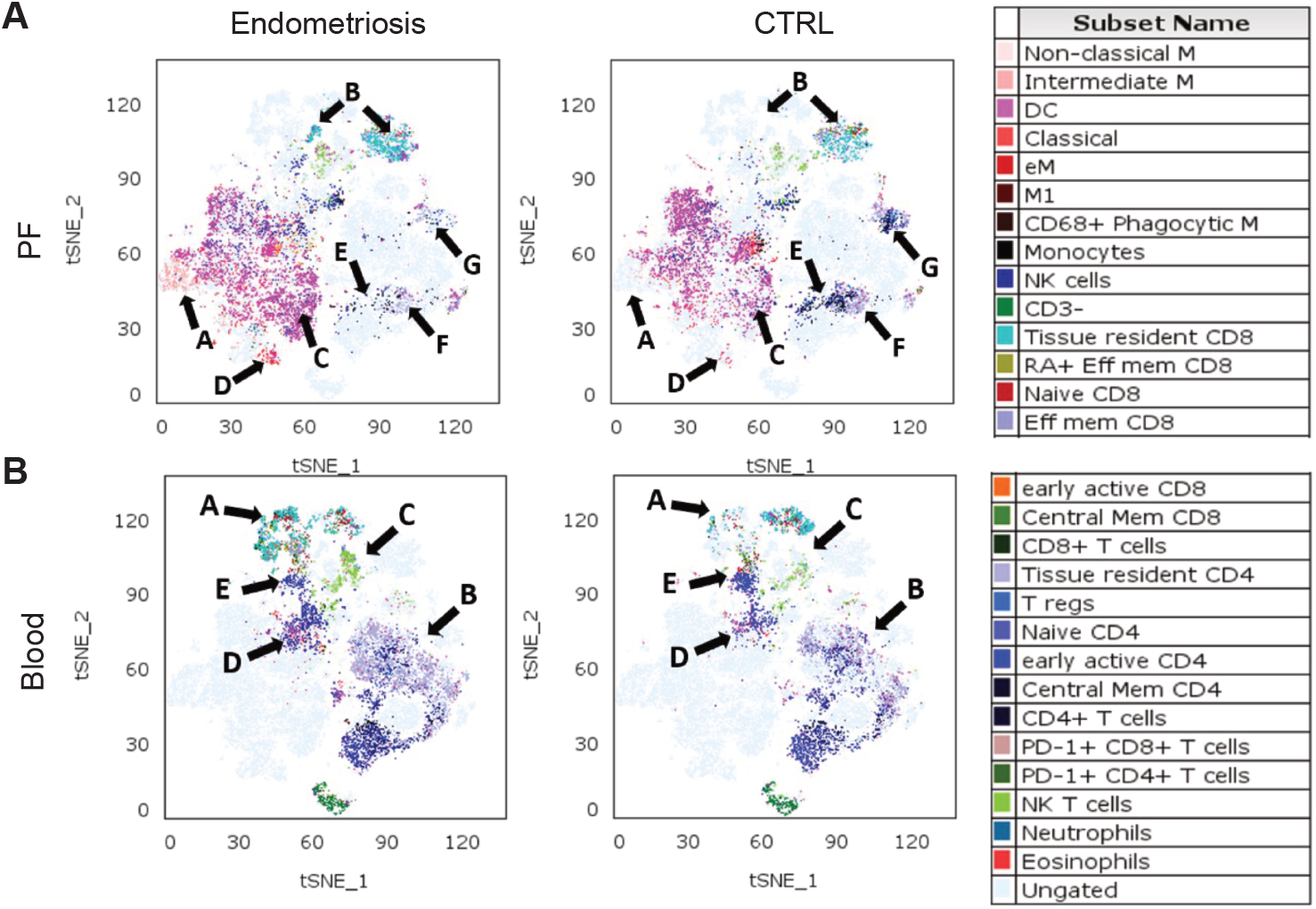
The immune environment varies demonstrably between peritoneal fluid (PF) and blood, and between endometriosis patients and controls. Blood and peritoneal fluid (PF) samples from endometriosis patients and controls were analysed by full-spectrum flow cytometry, concatenated, displayed using the viSNE algorithm^32^ and separated according to sample species (A, peritoneal fluid cells, n=5 endometriosis and n=4 control samples; B, blood cells, n=4 endometriosis and n=3 control samples) and disease state. The resulting maps of immune cell populations show marked differences in the myeloid and lymphoid compartments between endometriosis and control groups. Highlighted populations in PF (A): A, intermediate monocytes, B, CD8^+^ T cells, C, Dendritic Cells, D, endometrial macrophages, E, CD4^+^ T cells, F, CD4^+^ T_RM_ cells, G, CD4^+^ T_CM_ cells. In blood (B): A, CD4^+^ T_RM_ cells, B, CD4^+^ T cells, C, NKT cells, D, Dendritic Cells, E, early active (EA) CD4^+^ T cells. Data were analysed using FlowJo.

In peritoneal fluid, we found intermediate monocytes (Fig.1 A, population (pop) A) in the endometriosis samples, a population absent in the control group. Additionally, the endometriosis group exhibited a higher prevalence of CD8^+^ T cells (pop B), along with increased numbers of DCs (pop C) and endometrial macrophages (Mϕ, pop D) compared to controls. Conversely, the control group showed a higher prevalence of CD4^+^ T cells (pop E), CD4^+^ T_RM_ cells (pop F), and CD4^+^ T_CM_ cells (pop G).

In blood, the picture was almost opposite (Fig. 1 B): Within the endometriosis group, we observed a broader distribution of CD8^+^ T_RM_ cells (pop A) and CD4^+^ T cells (pop B), as well as an increase in NK T cells (pop C) and DCs (pop D). On the other hand, the control group showed an increase in early active CD4^+^ T cells (pop E).

To follow up these variations through the hierarchy of immune cell populations, we used automated clustering by x-shift and FlowSOM algorithms (Fig. 2). The distribution of the 14 populations allowed us to identify notable differences between endometriosis patients and controls. In peritoneal fluid, the endometriosis group exhibited higher percentages of DCs (pop 1), intermediate monocytes (pop 0), CD68^+^ Mϕ (pop 2), and CD8^+^ T cells (pop 8) compared to the control group. Specifically, CD68^+^ Mϕ constituted 31.6% in endometriosis patients and 12.9% in controls, DCs were 25.9% in endometriosis and 0.37% in controls, and intermediate monocytes were 7.03% in endometriosis and 0.01% in controls (Supplementary Fig. 2). CD8^+^ T cells accounted for 14.4% in endometriosis versus 0.91% in controls. Conversely, the control group displayed higher percentages of CD4^+^ T cells (pop 9 and 10), monocytes (pop 3), and B cells (pop 4 and 6) than the patients’ group. Notably, NK cells were 13.83% in endometriosis and 0.71% in controls, while monocytes were 46.2% in controls and 0.13% in endometriosis (E).

**Figure 2:**
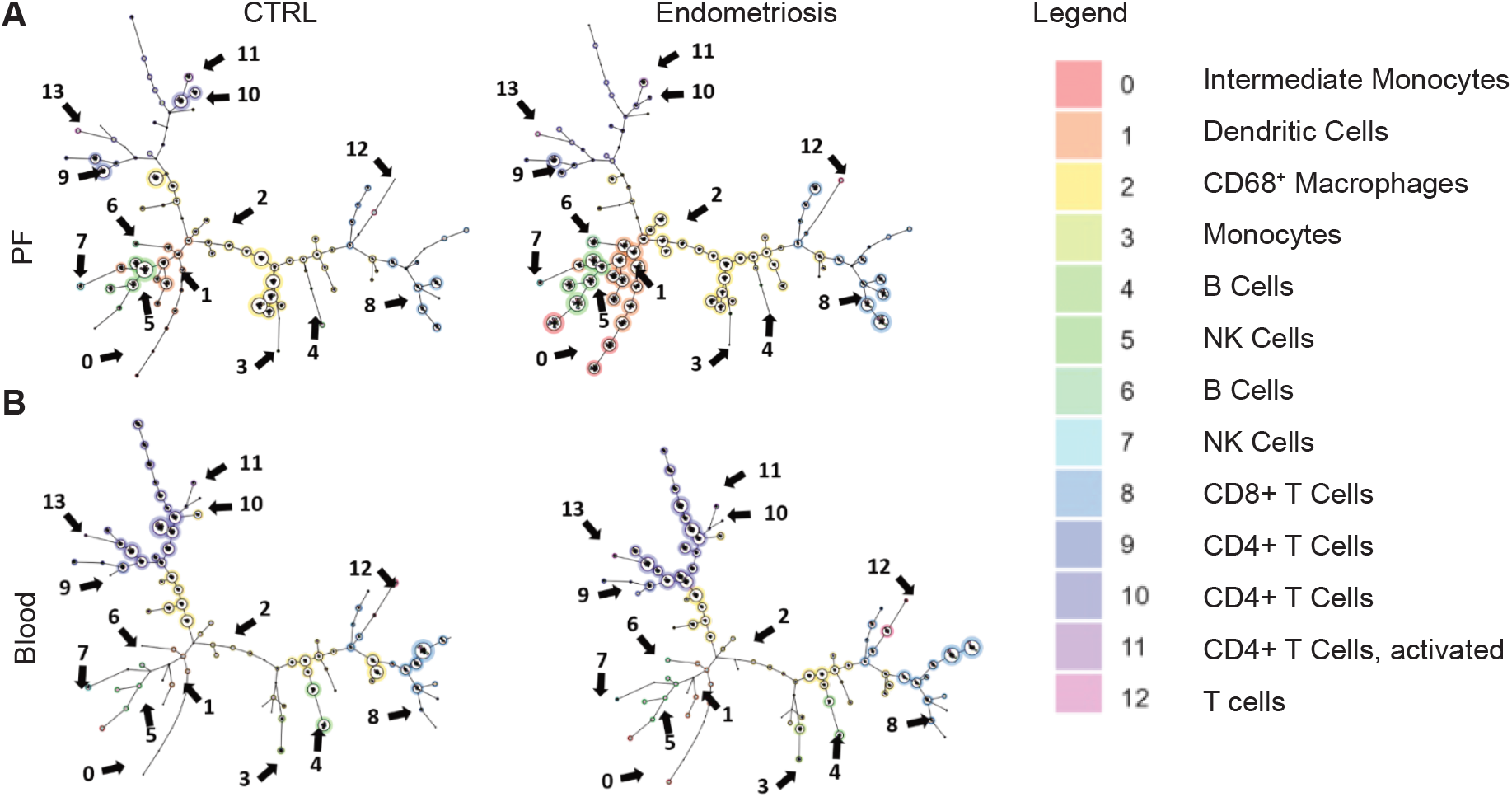
Monocyte and T cell subsets differ in endometriosis patients both in blood and peritoneal fluid. Blood and peritoneal fluid (PF) samples from endometriosis patients and controls were analysed by full-spectrum flow cytometry and the differences in hierarchic population sizes displayed using the x-shift and FlowSOM algorithms (A, peritoneal fluid cells, n=5 endometriosis and n=4 control samples; B, blood cells, n=4 endometriosis and n=3 control samples). Each node represents a cell cluster, its size the abundance of the population. The resulting minimum spanning tree graphs highlight the increase in the peritoneal myeloid compartment in endometriosis (A) and in the lymphoid compartment in the blood (B). Data were analysed using FlowJo.

Interestingly, in blood, endometriosis samples were characterised by having increased levels of CD8^+^ T cells, specifically in CD8^+^ T_RM_ cells and in Mϕ, specifically in CD68^+^ Mϕ (Fig. 2 B). However, investigating the differences between endometriosis patients and controls in blood samples was challenging due to the small sample size.

With regards to the role of immune cells in endometriosis pathophysiology, we next analysed the activation status of the respective populations in peritoneal fluid and blood cells by measuring the differences in expression of CD69, CD127, CD27, CD25, CD16, and CD40 between endometriosis and control populations (Fig. 3). In both, peritoneal fluid (Fig. 3 A) and blood (Fig. 3 B), the expression patterns of activation markers remained consistent between endometriosis patients and controls across the cell populations. In peritoneal fluid, activation was predominantly seen as an increase in CD69^+^ cells within the T cell compartment, an increase in CD16^+^ cells in the B cell and NK cell compartment, and an increase in CD40-expressing cells in the monocyte and DC compartment. In blood, the activation was a lot less prominent, with an increase only seen in CD16^+^ monocytes and dendritic cells.

**Figure 3:**
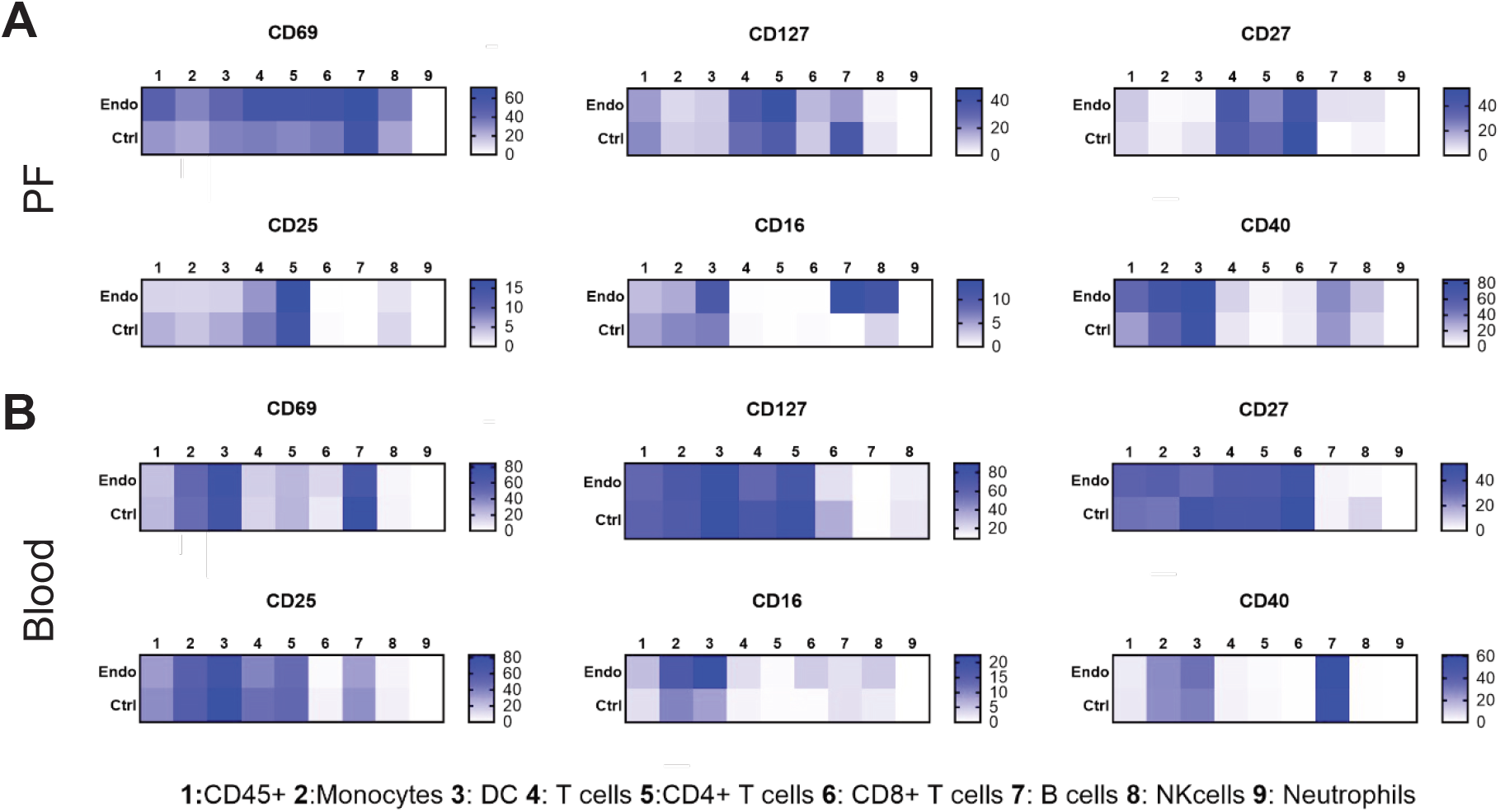
Activation of immune cells is tightly regulated in blood and more apparent in peritoneal fluid. Heatmaps showing the expression of activation markers on cells of various populations (1—9) in peritoneal fluid (A) and blood (B) of endometriosis patients (n=5) and controls (n=4). The scale bar indicates the percentage of cells expressing the respective marker within the total of each population. Differences in activation are mainly seen in PF populations (CD69, CD16, CD40), with blood more tightly regulated. (The notable exception is CD16.)

The active T cell compartment in PF led us to wonder whether an increased expression of immune regulatory mechanisms or immune checkpoints could explain the persistence of endometriosis lesions within the peritoneal cavity. We thus investigated the expression of FOXP3 as an indicator of T_REG_ presence as well as the expression of PD-1 as the paradigmatic immune checkpoint in PF and blood cells (Fig. 4). While no significant differences were seen with regards to FOXP expression, we were surprised to find a significant decrease in PD-1 expression overall in the PF cells (Fig. 4 A), and an inverse picture in blood (Fig. 4 B) – although our sample numbers were too low to reach significance here.

**Figure 4:**
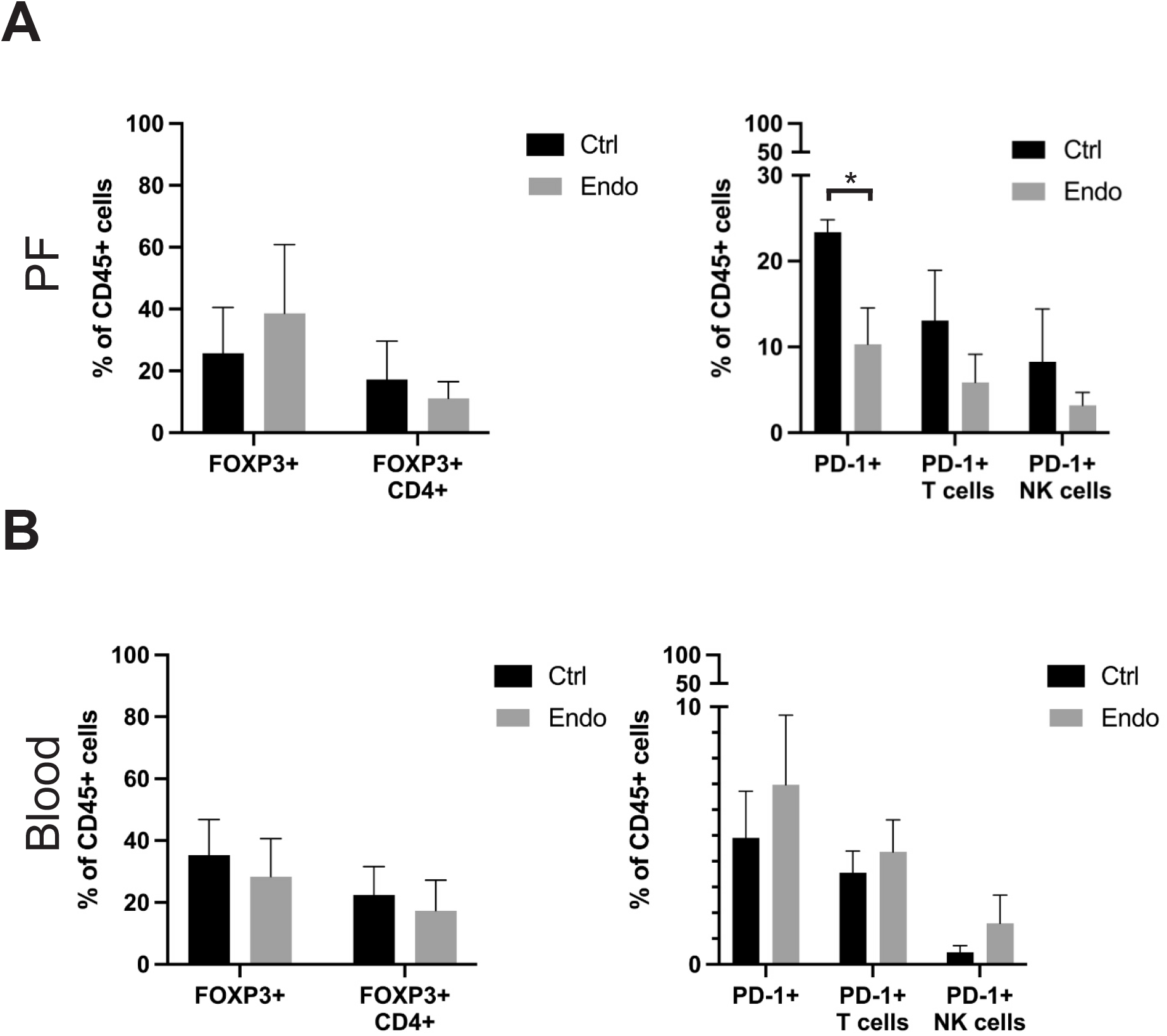
The immune checkpoint inhibition through PD-1 is diminished in the peritoneal fluid in endometriosis. The overall expression of T_REG_ marker FOXP3 and the immune checkpoint marker PD-1 was measured on overall CD45^+^ cells and subpopulations in peritoneal fluid (A) and blood (B) cells from endometriosis patients (n=5) and controls (n=4). *, p<0.05, Mann-Whitney U-test.

Overall, our results indicate an over-activated state of the lymphoid compartment, i.e., PD-1^+^CD4^+^ T cells, in the peritoneal fluid of women with endometriosis, which likely propagates the inflammation. Increasing the expression of PD-1 in relevant T cell clones could therefore be a new avenue of treating endometriosis-related inflammation and pain.

## Discussion

### Principal Findings

Here, we investigated the composition, activation status and immunoregulation of peritoneal fluid cells and blood in women with and without endometriosis. We found the PF cells dominated by the myeloid compartment and the blood by the lymphoid compartment, respectively; however, the main differences in activation and regulation between patients and controls were in the lymphoid compartment in PF and in the myeloid compartment in blood. While we had expected so see increased regulation and inhibition of PF T cells in endometriosis, surprisingly, the opposite pattern emerged, with the immune checkpoint marker PD-1 significantly decreased.

### Results in the Context of What is Known

Our results point towards a prominent role for PD-1 dependent inhibition of T cell immunity than previously thought. Earlier studies found an increase in activated CD69^+^ T cells in the peritoneal fluid of women with endometriosis^25^; we now add the interesting observation that PD-1 levels are decreased in this environment, suggesting an explanation for the increased activation seen earlier.

### Clinical Implications

The observed over-activation of T cells within the peritoneal fluid would—if confirmed—open an interesting angle of anti-inflammatory therapy in endometriosis.

### Research Implications

The question of how and why endometriosis lesions persist in the peritoneal cavity remains unanswered. It would also be interesting to investigate whether the T cells lacking PD-1 expressions are clonal, and what antigen they might recognise in endometriosis.

### Strengths and Limitations

A strength of this study is the use of full-spectrum flow cytometry to analyse immune cell populations in depth in both, blood and peritoneal fluid. This has previously been done by us^25^ and others^31^ using cytometry by time-of-flight (CyTOF); however, FSFC offers a broader range of antibodies to use and allows for easier translation of existing marker panels into high-dimensional assays.

A clear weakness of our study is the low sample number – with only 9 PF and 7 blood samples, any conclusions will have to be carefully drawn. We also were unable to achieve significance in many assessments even if they looked to indicate a clear trend.

## Conclusions

The dysregulation of the immune system in endometriosis, and especially the lack of intrinsic inhibition offers new avenues of therapy. The question of whether the over-activated T cell compartment is clonal in nature is exciting and warrants further investigation.

## Supporting information

Suppl Figure 1 Gating

Suppl Table 2 Antibodies

Suppl Table 3 Cell Populations

Suppl Table 2 Participants

Suppl Figure 2 All Cells Panel

