## Supplementary material for "The over-activated peritoneal immune environment in endometriosis is characterised by a lack of PD-1 inhibition": Suppl Figure 1 Gating

Supplementary Figure Gating

Myeloid compartment (granulocytes and monocytes)

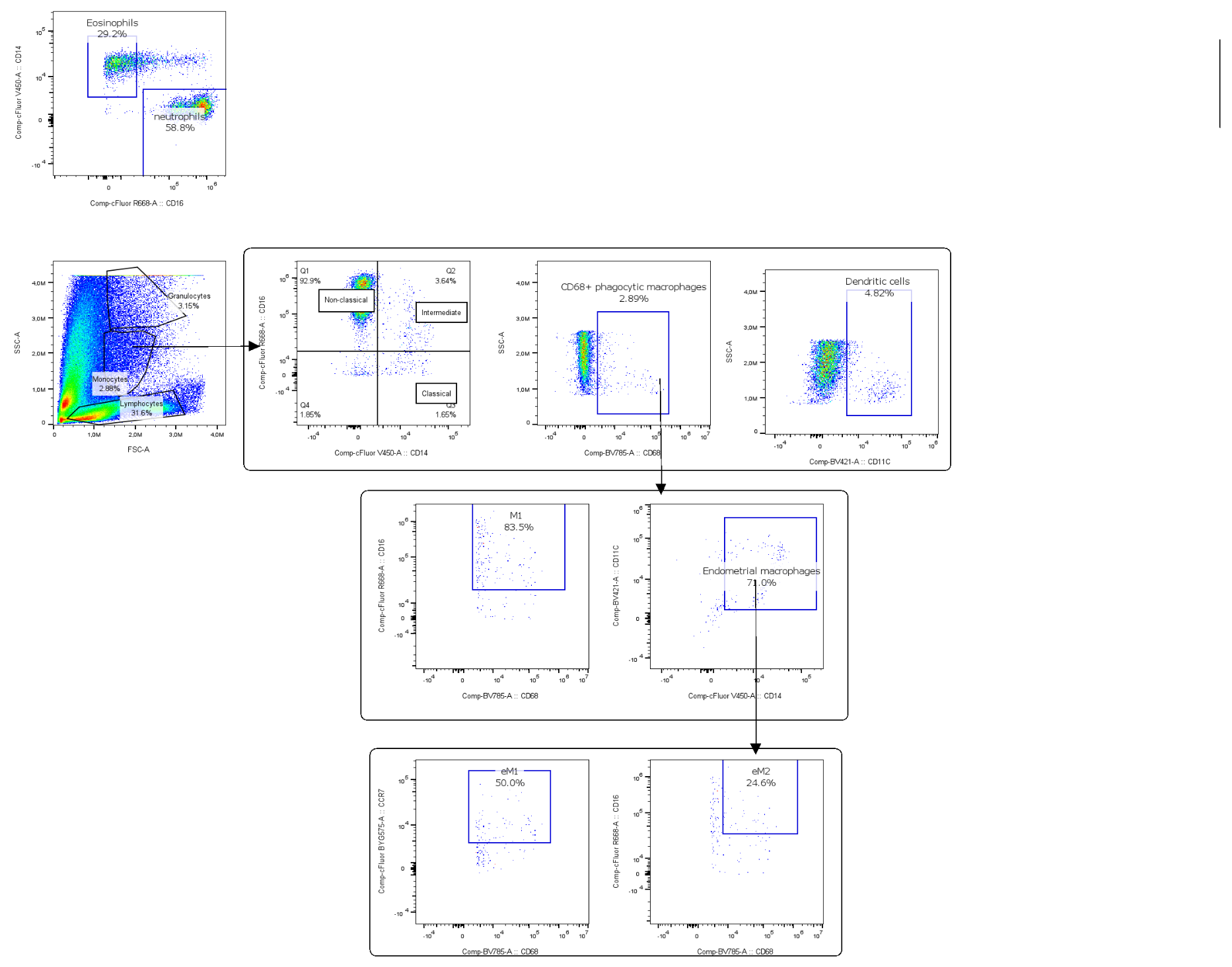

Lymphoid compartment (T cells, B cells, etc.)

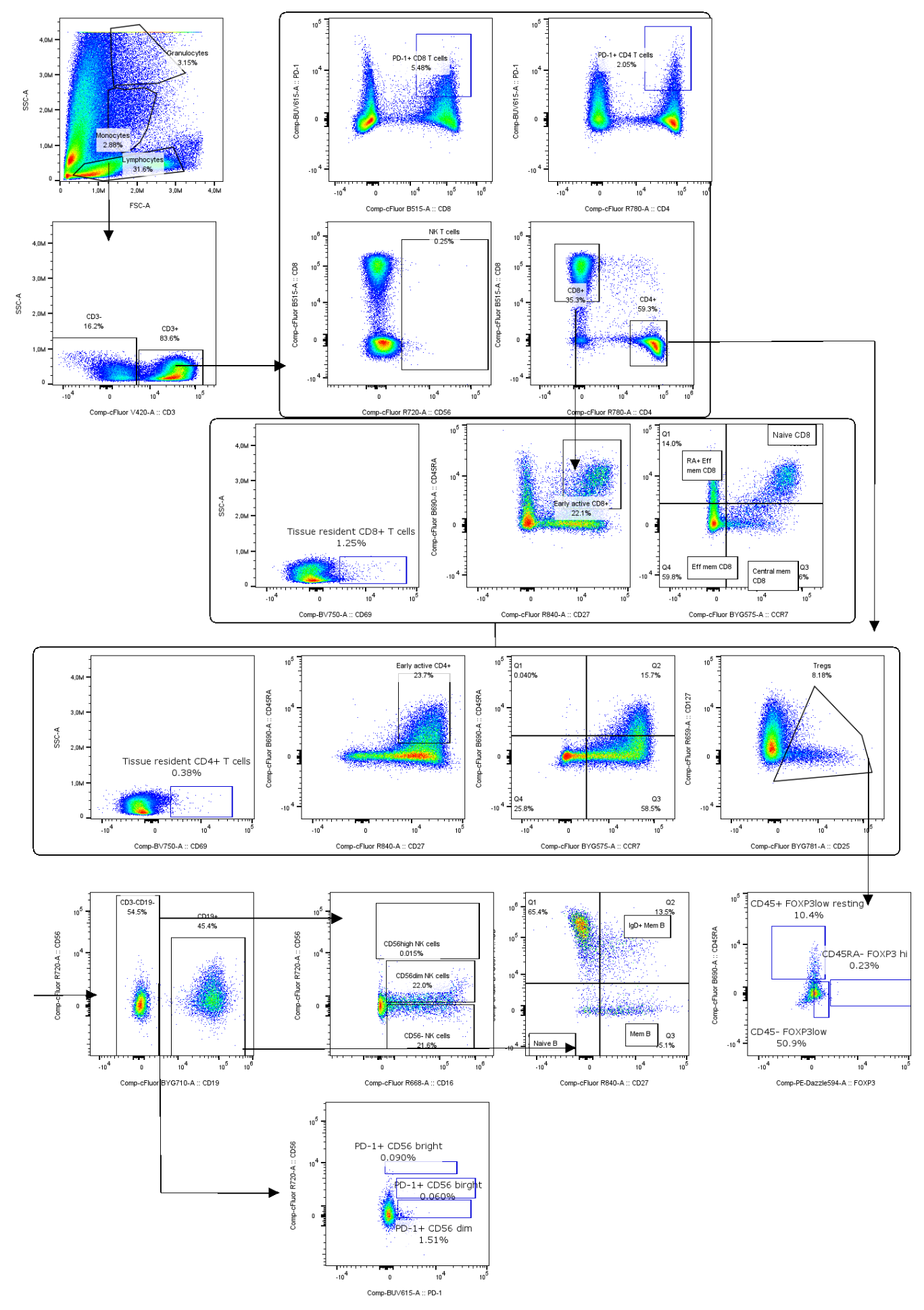
