## Supplementary material for "The over-activated peritoneal immune environment in endometriosis is characterised by a lack of PD-1 inhibition": Suppl Table 2 Antibodies

Table antibodies

| **Target** | **Clone** | **Fluorochrome** | **Manufacturer** | **Catalog  number** |
| --- | --- | --- | --- | --- |
| CD3 | SK7 | cFluor V420 | Cytek Biosciences | R7-20054 |
| CD4 | SK3 | cFluor R780 | Cytek Biosciences | R7-20084 |
| CD8 | SK1 | cFluor B515 | Cytek Biosciences | R7-20036 |
| CD11c | 3.9 | BV421 | BioLegend | 301628 |
| CD14 | M5E2 | cFluor V450 | Cytek Biosciences | R7-20004 |
| CD16 | 3G8 | cFluor R668 | Cytek Biosciences | R7-20070 |
| CD19 | HIB19 | cFluor BYG710 | Cytek Biosciences | R7-20010 |
| CD25 | BC96 | cFluor BYG781 | Cytek Biosciences | R7-20080 |
| CD27 | QA17A18 | cFluor R840 | Cytek Biosciences | R7-20082 |
| CD40 | 5C3 | PE-Fire 810 | BioLegend | 334355 |
| CD45 | HI30 | cFluor V547 | Cytek Biosciences | R7-20012 |
| CD45RA | HI100 | cFluor B690 | Cytek Biosciences | R7-20086 |
| CD56 | 5.1H11 | cFluor R720 | Cytek Biosciences | R7-20090 |
| CD68 | Y1/82A | BV785 | BioLegend | 3333826 |
| CD69 | FN50 | BV750 | BioLegend | 310954 |
| CD127 | A019D5 | cFluor R659 | Cytek Biosciences | R7-20078 |
| CD197 (CCR7) | G043H7 | cFluor BYG575 | Cytek Biosciences | R7-20076 |
| CD279 (PD-1) | EH12.1 | BUV615 | BD Biosciences | 612991 |
| IgD | IA6-2 | cFluor BYG667 | Cytek Biosciences | R7-20088 |
| FOXP3 | 206D | PE-Dazzle 594 | BioLegend | 320126 |
| VIABILITY |  | ViaDye™ Red | Cytek Biosciences | R7-60008 |
