## Supplementary material for "The over-activated peritoneal immune environment in endometriosis is characterised by a lack of PD-1 inhibition": Suppl Table 3 Cell Populations

Table Cell Populations

| **Cells** | **Type** | **Subtype** | **Markers** |
| --- | --- | --- | --- |
| **Hematopoietic cells** |  | | Viadye-CD45^+^ |
| **Monocytes** |  | | FSC-A SSC-A |
|  | Peripheral monocytes |  | |
|  |  | Non-classical | CD16^+^ |
|  |  | Intermediate | CD14^+^ CD16^+^ |
|  |  | Classical | CD14^+^ |
|  | Peripheral Mϕ |  | CD68^+^ SSC-A |
|  |  | M1 | CD16^+^ |
|  | Endometrial Mϕ |  | CD68^+^ |
|  |  |  | CD14^+^ CD11c^+^ |
|  |  | eM1 | CCR7^+^ |
|  |  | eM2 | CD16^+^ |
|  | Dendritic cells |  | CD11c^+^ SSC-A |
| **Lymphocytes** |  | | FSC-A SSC-A |
| **T cells** |  | | CD3^+^ |
|  | PD-1 expressing T cells |  | |
|  |  | CD4 | CD4^+^ PD-1^+^ |
|  |  | CD8 | CD8^+^ PD-1^+^ |
|  | CD4 T cells |  | CD4^+^ |
|  |  | Early active CD4 T cells | CD27^+^CD45RA^+^ |
|  |  | Naïve CD4 T cells | CCR7^+^CD45RA^+^ |
|  |  | Central memory CD4 T cells | CCR7^+^CD45RA^-^ |
|  |  | Effector memory CD4 T cells | CCR7^-^CD45RA^-^ |
|  |  | Tissue resident T cells | CD69^+^ SSC-A |
|  | CD8 T cells |  | CD8^+^ |
|  |  | Early active CD8 T cells | CD27^+^ CD45RA^+^ |
|  |  | Naïve CD8 T cells | CCR7^+^ CD45RA^+^ |
|  |  | Central memory CD8 T cells | CCR7^+^CD45RA- |
|  |  | Effector memory CD8 T cells | CCR7^-^CD45RA- |
|  |  | RA+ effector memory CD8 T cells | CCR7^-^CD45RA |
|  |  | Tissue resident T cells | CD69^+^ SSC-A |
|  | Regulatory T cells |  | CD25^+^ CD127^+^ |
|  |  | Resting | CD45RA^+^ FOXP3 LOW |
|  |  | Activated | CD45RA^-^ FOXP3 ^HI^ |
|  |  | Non suppressive | CD45RA^-^FOXP3 ^LOW^ |
|  | NK T cells |  | CD56^+^CD8^+^ |
| **NK cells** |  | | CD3^-^CD19^-^ |
|  | Peripheral NK cells |  |  |
|  |  | CD56^-^ NK cells | CD56^-^ CD16+ |
|  |  | Cytotoxic NK cells | CD56^dim^ CD16^+^ |
|  |  | Regulatory NK cells | CD56^bright^ PD-1^+^ |
|  | Endometrial/uterine NK cells |  | CD56^superbright^ PD-1^+^ |
| **B cells** |  |  | CD19^+^ |
|  | Naïve B cells |  | IgD^+^ CD27- |
|  | IgD^+^ memory B cells |  | IgD^+^CD27^+^ |
|  | Memory B cells |  | IgD^-^ CD27^+^ |
| **Granulocytes** |  |  | FSC-A SSC-A |
|  | Neutrophils |  | CD16^+^ |
|  | Eosinophils |  | CD14^+^ |
| **Non hematopoietic cells** |  | | Viadye ^-^ CD45^-^ |
