## Supplementary material for "The over-activated peritoneal immune environment in endometriosis is characterised by a lack of PD-1 inhibition": Suppl Table 2 Participants

Table Participants

| **Sample ID** | **Age** | **Menstrual  Phase** | **Cause for surgery** | **Endo  diagnosis** | **ASRM stage** | **Group** |
| --- | --- | --- | --- | --- | --- | --- |
| 18/23 | 44 | Secretory | Hysterectomy Bilateral  salpingectomy USL suspension Colposuspension Posterior repair Cystoscopy | No | 0 | Control |
| 67/23 | 30 | Proliferative | Infertility Pelvic pain Ovulation pain | Yes | 1 | Patient |
| 95/23 | 25 | Secretory | Dysmenorrhea Menorrhegia | Yes | 1 | Patient |
| 98/23 | 35 | Secretory | Pelvic pain Adhesions | No | 0 | Control |
| 102/23 | 30 | Proliferative | Possible  endometriosis | Yes | 2 | Patient |
| 106/23 | 35 | Secretory | Dysmenorrhoea Endometriosis | Yes | 2 | Patient |
| 109/23 | 38 | Menses | Bilateral ovarian cysts Pelvic pain  Menorrhagia  Subfertility | No | 0 | Control |
| 115/23 | 27 | Secretory | Pelvic pain Dysuria | No | 0 | Control |
| 122/23 | 32 | Secretory | Mild to moderate  endometriosis  on USS | Yes | 1 | Patient |
