## Supplementary figures and images for "The over-activated peritoneal immune environment in endometriosis is characterised by a lack of PD-1 inhibition"

### Suppl Figure 2 All Cells Panel

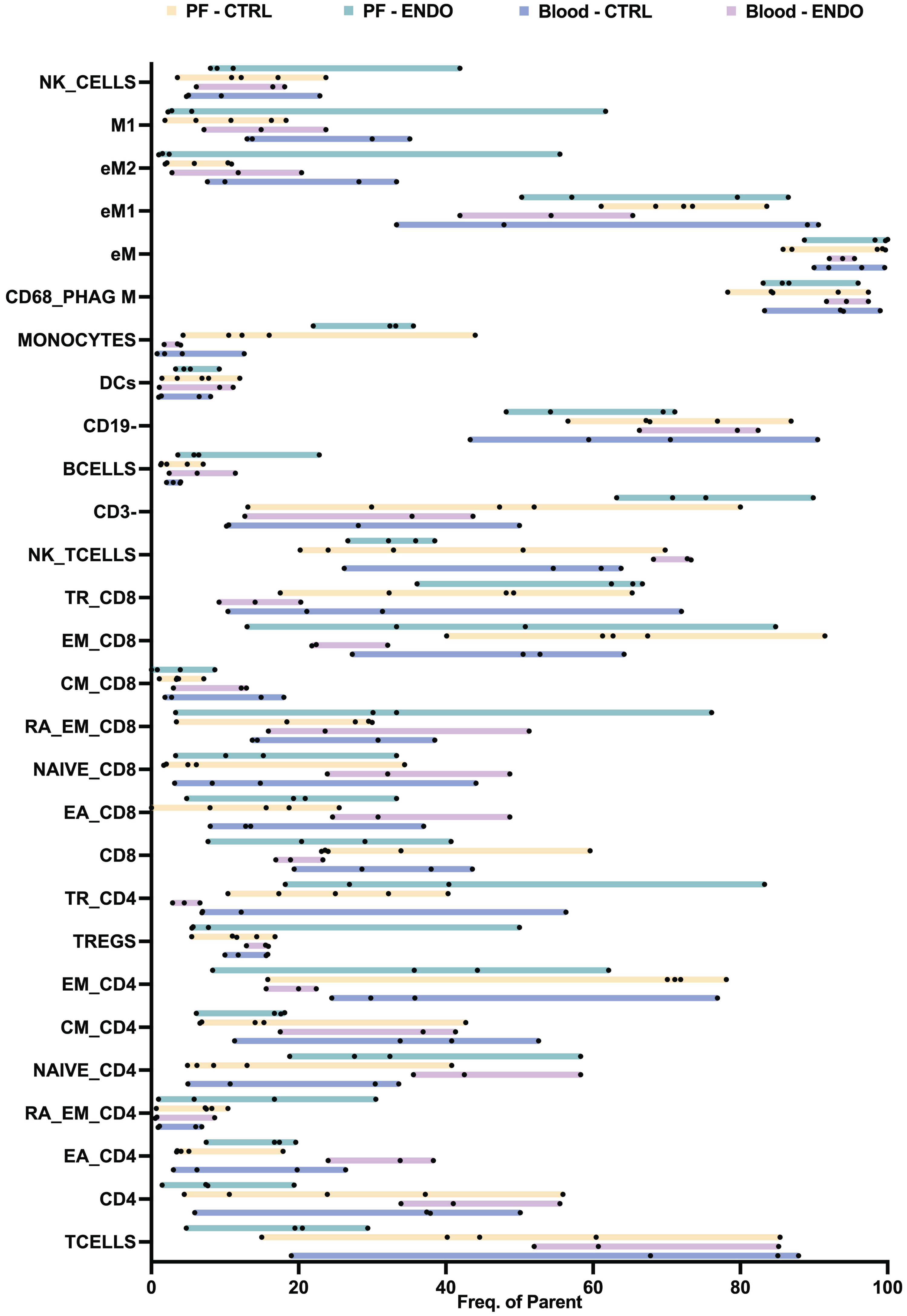
